# A Model For Sea Lice (*Lepeophtheirus salmonis*) Dynamics In A Seasonally Changing Environment

**DOI:** 10.1101/026583

**Authors:** Matthew A. Rittenhouse, Crawford W. Revie, Amy Hurford

## Abstract

Sea lice (Lepeophtheirus salmonis) are a significant source of monetary losses on salmon farms. Sea lice exhibit temperature-dependent development rates and salinity-dependent mortality, but to date no deterministic models have incorporated these seasonally varying factors. To understand how environmental variation and life history characteristics affect sea lice abundance, we derive a delay differential equation model and parameterize the model with environmental data from British Columbia and southern Newfoundland. We calculate the lifetime reproductive output for female sea lice maturing to adulthood at different times of the year and find differences in the timing of peak reproduction between the two regions. Using a sensitivity analysis, we find that sea lice abundance is more sensitive to variation in mean annual water temperature and mean annual salinity than to variation in life history parameters. Our results suggest that effective sea lice management requires consideration of site-specific temperature and salinity patterns and, in particular, that the optimal timing of production cycles and sea lice treatments might vary between regions.

## 1 Introduction

The control of parasitic organisms is a major concern in marine aquaculture. In particular, sea lice *(Lepeophtheirus salmonis* and *Caligus spp.)* cause substantial economic losses on salmon farms (Costello 2009). Due to their economic importance, control of sea lice on salmon farms has been named one of the top priorities in aquaculture research by both scientists and aquaculture practitioners (Jones et al. 2014). Adequate control of sea lice is predicated on the ability to predict future lice levels from current population and environmental trends, as well as predicting the effectiveness of different treatment regimes. These two needs can be accomplished through mathematical modelling and it is imperative that tractable and biologically sound models are developed to aid practitioners in decisions regarding sea lice dynamics.

Seasonal environmental variability plays a major role in the dynamics of many disease systems (Altizer et al. 2006). Temperature and salinity affect several characteristics of sea lice life history (summarized in Table 1), thus models of sea lice dynamics must be able to incorporate the effects of seasonally varying temperature and salinity on the sea louse lifecycle.

**Table 1:**
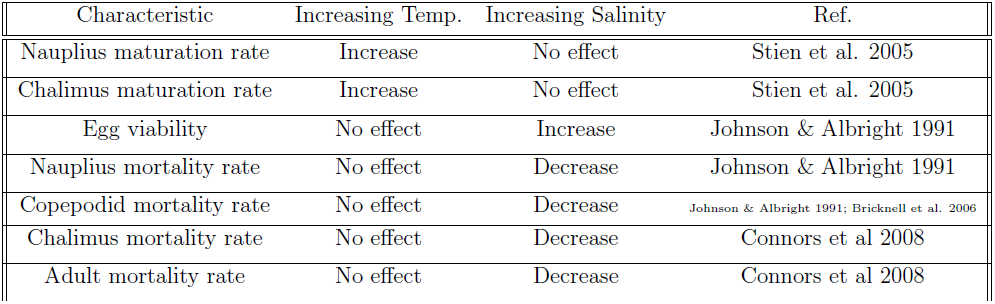
The effect of water temperature and salinity on characteristics of sea lice life history.

| Characteristic | Increasing Temp. | Increasing Salinity | Ref. |
| --- | --- | --- | --- |
| Nauplius maturation rate | Increase | No effect | Stien et al. 2005 |
| Chalimus maturation rate | Increase | No effect | Stien et al. 2005 |
| Egg viability | No effect | Increase | Johnson & Albright 1991 |
| Nauplius mortality rate | No effect | Decrease | Johnson & Albright 1991 |
| Copepodid mortality rate | No effect | Decrease | Johnson & Albright 1991; Bricknell et al. 2006 |
| Chalimus mortality rate | No effect | Decrease | Connors et al 2008 |
| Adult mortality rate | No effect | Decrease | Connors et al 2008 |

A variety of deterministic (Revie et al 2005, Stien et al. 2005, Robbins et al. 2010, Gettinby et al. 2011, Aldrin et al. 2013, Groner et al. 2014, Kristoffersen et al. 2014) and stochastic (Aldrin et al. 2013, Groner et al. 2014) models have been derived to predict sea lice dynamics. Revie et al. (2005) derived a life stage model with fixed delays and constant mortality rates that formed the basis for the simulation tool, SLiDESim. Robbins et al. (2010) utilized SLiDESim to search for optimal treatment strategies in Scottish farms. Gettinby et al. (2011) tested the SLiDESim model on sea lice collection data from the Hardangerfjord in south-west Norway. These authors concluded that for the model to be utilized in evaluating treatment strategies, a better understanding of the underlying biological and environmental factors, including temperature-dependent maturation and salinity-dependent survival, was necessary. Kristoffersen et al. (2014) used Bĕlehrádek functions derived in Stien et al. (2005) to estimate degree-days needed for sea lice to mature from one stage to the next and derived a life stage model with temperature-based delays and constant mortality rates. Aldrin et al (2013) used a stochastic spatio-temporal model to show how seawater temperatures, fish stock population, and distance between farms contributed to predicted sea lice counts. Groner et al. (2014) created a stochastic matrix population model to examine the effects of seasonally varying temperature on treatment schemes and louse mate limitation.

A deterministic model capable of accounting for both the effects of seasonally varying temperature on sea lice maturation, and the effects of seasonally varying salinity on sea lice mortality has not yet been developed. Stien et al. 2005 suggest that the delay differential equation models described in Nisbet & Gurney (1983) as a method to address this need. Delay differential equations models of this type have been successfully used in epidemiological models of koi herpes virus (Omori & Adams 2010) and malaria (Beck-Johnson et al. 2014).

We present a delay differential equation model of the sea lice lifecycle with temperature-dependent stage durations, salinity-dependent mortality, and time-dependent temperature/salinity. Where possible, model parameters are fitted to values from the literature for the species, *Lepeophtheirus salmonis.* The time dependent basic reproductive ratio, *R*_0_ (*t*), is derived numerically to quantify seasonal differences in sea lice replenishment at sites in British Columbia and southern Newfoundland, Canada. Additionally, a sensitivity analysis is conducted to identify the parameters that most substantially affect the model predictions.

## 2 The Model

We developed a model for sea lice dynamics on salmon farms that includes temperature-dependent maturation delays and salinity-dependent mortality. *Lepeophtheirus salmonis* exhibit 8 distinct life stages, consisting of nauplius I/II, copepodids, chalimus I/II, pre-adult I/II, and adults (Hamre et al 2013). For the purpose of modelling, we assume that sea lice may be in 1 of 4 possible functional states: planktonic non-infectious nauplii (*P*), infectious copepodids (*I*), non-reproductive chalimus and pre-adults (*C*), or adult females, (*A*). Each individual matures through the states in order from nauplius (*P*), to copepodid, (*I*), to chalimus (*C*), and finally to adult female (*A*) (Fig. 1).

**Figure 1:**
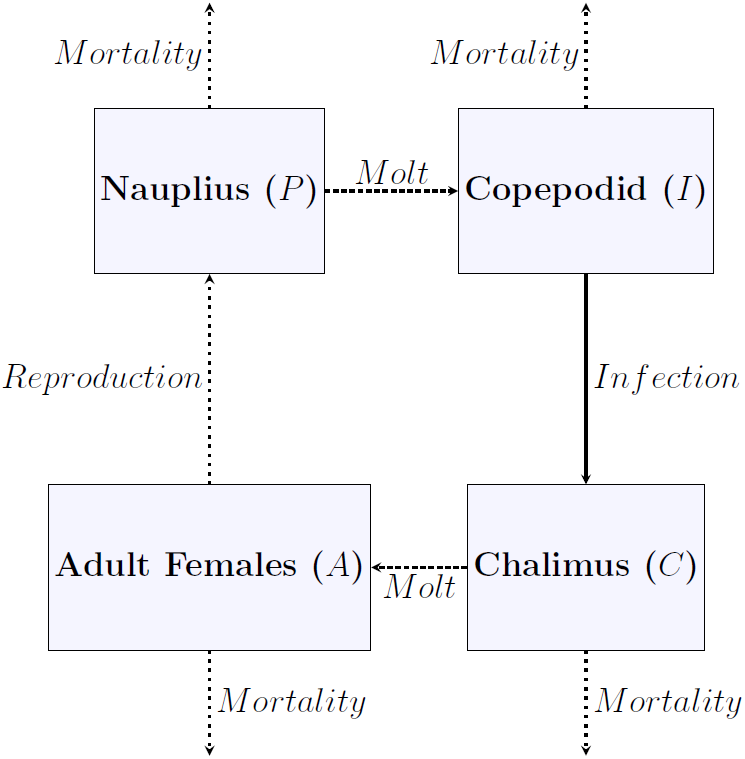
Modelled life cycle of the sea louse. Dashed arrows indicate aspects of the life history affected by temperature. Dotted arrows indicated aspects of the life history affected by salinity.

The length of time that a nauplius or chalimus requires to mature to their respective next life stages depends on water temperature (Table 1). Let *γ*_*x*_(*T*(*t*)) be a function that describes the rate of change in the level of development for a given stage *x* ∈ {*P,C*} as it depends on temperature (*T*), which changes over time (*t*). For notational simplicity, we write simply *γ*_*x*_(*t*), because given functions that describe how temperature changes with respect to time (*T*(*t*)), and how the development rate changes with respect to temperature (*γ*_*x*_(*T*)), we can then determine how the development rate changes with respect to time (*γ*_*x*_(*t*)) without needing to explicitly reference the dependence on temperature.

The waiting times associated with maturation are such that a cohort exiting a state *x* at time *t*, will all have entered that stage at *t* − *τ*_*x*_(*t*). The waiting time, *τ*_*x*_(*t*), depends on the development rate, *γ*_*x*_(*t*), and is defined as the length of time that it takes sea lice to reach a threshold development level, 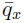, given that they entered the stage *x* with a development level *q*_*x*_ = 0. As such, *τ*_*x*_(*t*) is implicitly defined as,

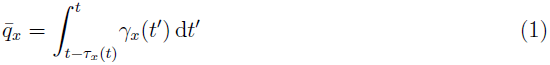

(Nisbet & Gurney, 1983).

Natural mortality occurs in all stages at a per capita rate *μ*_*y*_(*S*(*t*)), where *y* ∈ {*P, I, C, A*}. Natural mortality is a function of salinity *S*(*t*), which is a function of time (*t*). For notational simplicity, we write simply *μ*_*y*_(*t*), because given functions that describe how salinity changes with respect to time (*S*(*t*)), and how the mortality rate changes with respect to salinity *(μ_y_*(*S*)), we can then determine how the mortality rate changes with respect to time (*μ*_*y*_(*t*)) without needing to explicitly reference the dependence on salinity. Not all members of a cohort who enter a stage *x* at time *t* − *τ*_*x*_(*t*) survive to mature at time *t*. The proportion of the cohort that survive the maturation period is,

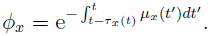

The proportion of eggs that produce viable nauplii is a function of salinity, *v*(*t*). All other events in the sea lice life history do not depend on temporally varying quantities and are assumed to occur at constant per capita rates. The complete model is a system of delay differential equations,

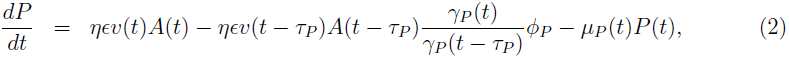

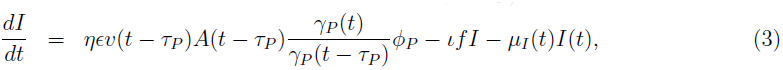

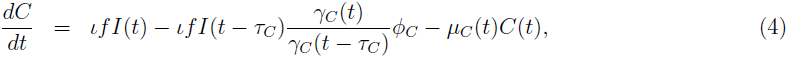

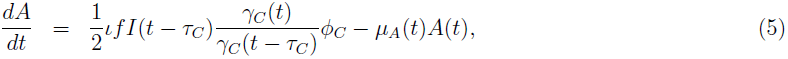

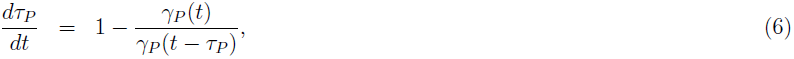

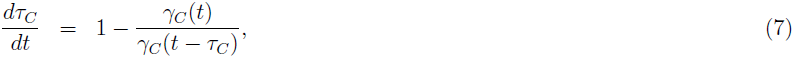

where *η* is the number of eggs per egg string, *ε* is the rate of egg string production, *ι* is the rate of infection per fish, *f* is the number of fish on the farm, and all model parameters are summarized in Table 2. Equations (6) and (7) arise from differentiating equation (1) with respect to time (Nisbet and Gurney, 1983). The *γ*_*x*_(t)/*γ*_*x*_(*t* − *τ*_*x*_) terms arise because we wanted to ensure a correspondence between our model (which lumps all individuals with a development level 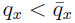 together into one state) and a model that treats the development level as a continuous quantity (Nisbet & Gurney, 1983; see Appendix A for further details).

**Table 2:**
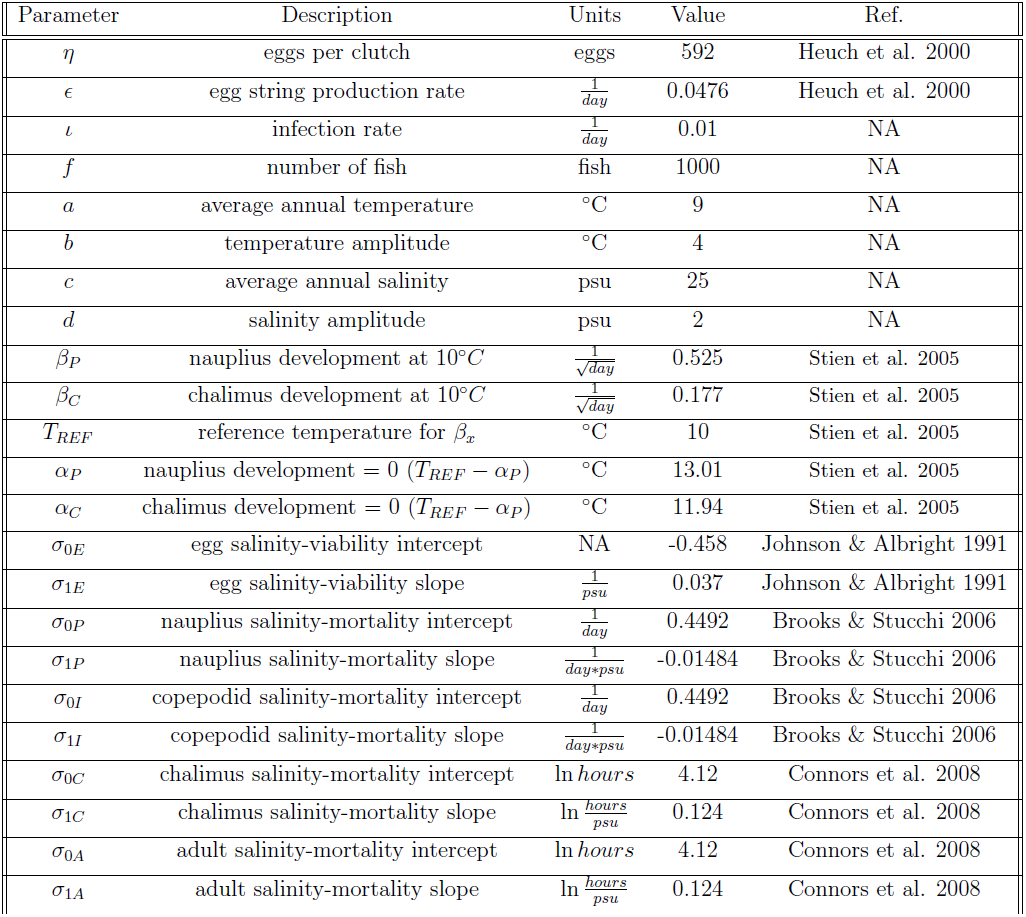
Model Parameters

| Parameter | Description | Units | Value | Ref. |
| --- | --- | --- | --- | --- |
| $\eta$ | eggs per clutch | eggs | 592 | Heuch et al. 2000 |
| $\epsilon$ | egg string production rate | $\frac{1}{day}$ | 0.0476 | Heuch et al. 2000 |
| $\iota$ | infection rate | $\frac{1}{day}$ | 0.01 | NA |
| $f$ | number of fish | fish | 1000 | NA |
| $a$ | average annual temperature | $^{\circ}C$ | 9 | NA |
| $b$ | temperature amplitude | $^{\circ}C$ | 4 | NA |
| $c$ | average annual salinity | psu | 25 | NA |
| $d$ | salinity amplitude | psu | 2 | NA |
| $\beta_P$ | nauplius development at $10^{\circ}C$ | $\frac{1}{\sqrt{day}}$ | 0.525 | Stien et al. 2005 |
| $\beta_C$ | chalmus development at $10^{\circ}C$ | $\frac{1}{\sqrt{day}}$ | 0.177 | Stien et al. 2005 |
| $T_{REF}$ | reference temperature for $\beta_x$ | $^{\circ}C$ | 10 | Stien et al. 2005 |
| $\alpha_P$ | nauplius development = 0 ($T_{REF} - \alpha_P$) | $^{\circ}C$ | 13.01 | Stien et al. 2005 |
| $\alpha_C$ | chalmus development = 0 ($T_{REF} - \alpha_P$) | $^{\circ}C$ | 11.94 | Stien et al. 2005 |
| $\sigma_{0E}$ | egg salinity-viability intercept | NA | -0.458 | Johnson & Albright 1991 |
| $\sigma_{1E}$ | egg salinity-viability slope | $\frac{1}{psu}$ | 0.037 | Johnson & Albright 1991 |
| $\sigma_{0P}$ | nauplius salinity-mortality intercept | $\frac{1}{day}$ | 0.4492 | Brooks & Stucchi 2006 |
| $\sigma_{1P}$ | nauplius salinity-mortality slope | $\frac{1}{day*psu}$ | -0.01484 | Brooks & Stucchi 2006 |
| $\sigma_{0I}$ | copepodid salinity-mortality intercept | $\frac{1}{day}$ | 0.4492 | Brooks & Stucchi 2006 |
| $\sigma_{1I}$ | copepodid salinity-mortality slope | $\frac{1}{day*psu}$ | -0.01484 | Brooks & Stucchi 2006 |
| $\sigma_{0C}$ | chalmus salinity-mortality intercept | $\ln hours$ | 4.12 | Connors et al. 2008 |
| $\sigma_{1C}$ | chalmus salinity-mortality slope | $\ln \frac{hours}{psu}$ | 0.124 | Connors et al. 2008 |
| $\sigma_{0A}$ | adult salinity-mortality intercept | $\ln hours$ | 4.12 | Connors et al. 2008 |
| $\sigma_{1A}$ | adult salinity-mortality slope | $\ln \frac{hours}{psu}$ | 0.124 | Connors et al. 2008 |

It should be noted that our model lacks a mechanism for density dependence in the sea lice population. Sea lice could exhibit density dependence due to an Allee effect caused by difficulties in mate finding at low densities (Krkošek et al. 2012, Groner et al. 2014). They may also exhibit density dependence due to host mortality or decreased reproduction at high intensities. In an aquaculture setting, managers will typically intervene with chemotherapeutics before any natural density-dependent regulation of the sea louse population can occur.

## 3 Model Parameterization

For our model, the maturation rate is the inverse of the Bĕlehrádek functions describing minimum development time as a function of temperature (Stien et al. 2005),

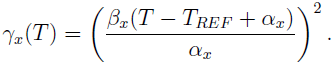

The shape of the function is described by the duration of the life stage 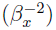 at the reference temperature *(T_REF_*) and by the location of an asymptote at *T*_*REF*_ − *α*_*x*_. The terms *β*_*P*_ and *β*_*C*_ are identical to the *β*_*2*_ terms for time from hatching to copepodid and time from infection to adult females in Stien et al. (2005). The *α*_*P*_ and *α*_*C*_ terms are the product of *β*_1_ and *β*_2_ from Stien et al. (2005). The reference temperature is 10°C.

Water temperature (T*(t)*) and salinity (S*(t)*) on salmon farms varies over time. We use sinusoidal functions to describe the general annual patterns,

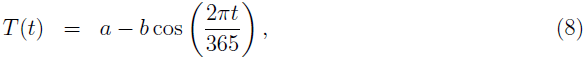

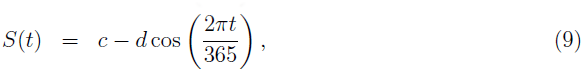

where *a* is the average annual temperature, *c* is the average annual salinity, and *b* and *d* are the respective amplitudes of the cosine functions. Sinusoidal functions of the form,

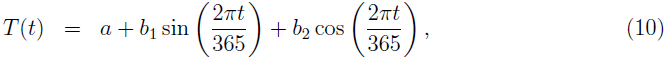

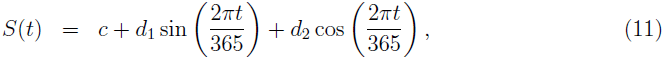

were fit to monthly temperature and salinity data from a salmon farm in the Broughton Archipelago of British Columbia (Marty et al., 2010; Fig. 4D and E) and to quarter-hourly temperature and salinity data from a salmon farm on the southern coast of Newfoundland (data provided by the Aquaculture Real-Time Integrated Environmental System; Fig. 4D and E).

Mortality is related to salinity via linear and log-linear relationships from the literature. The salinity-mortality relationship *μ*_*A*_(*S*)) is log-linear for adult sea lice (Connors et al. 2008). We assume that the mortality rate for adults and chalimi (*μ*_*C*_ (*S*)) is similar. The salinity-mortality relationship for nauplii (*μ*_*P*_ (*S*)) is a linear model from Brooks & Stucchi (2006), which is fit to data from Johnson & Albright (1991). We assume that the mortality rate for nauplii and copepodids (*μ*_*I*_(*S*)) is similar.

The number of eggs per egg string (*η*) was parameterized using data from Heuch et al. (2000). The lower bound on the egg string production rate (*ε*) is taken from Mustafa et al. (2000) and is used as the default egg string production rate. The upper bound for the egg string production rate, used in the uniform distribution for the sensitivity analysis, was taken from Heuch et al. (2000). Egg viability (*ν*(*S*)) consists of a linear model fit to salinity-survivorship data from Johnson & Albright (1991).

The infection success rate (*ι*) is dependent on a number of variables and is not well understood at the farm level. As such, a wide range of values, from 0.001 to 0.9, were used in the sensitivity analysis and a value of 0.01 was chosen as the default value.

## 4 Model Dynamics

We numerically solved the system of equations (2)–(7) using the PBSddesolve package in R. Due to the lack of density dependence our model produces either unbounded growth or extinction (Figs. 2 and 3). The abrupt changes in the trajectory slope shown in Fig. 2 indicate the beginning of successive generations. When the initial infection occurs on the coldest day of the year, the second cohort will take less days to mature to adulthood than the first, due to warming temperatures. Sites with higher average annual temperatures will take less time for cohorts to reach adulthood than sites with lower average annual temperatures (Fig. 2).

**Figure 2:**
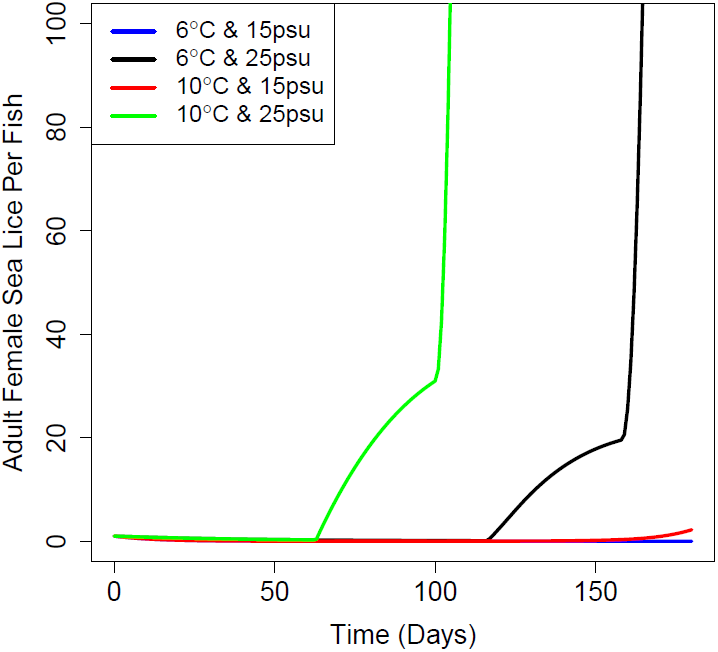
Four model simulations under high/high, high/low, low/high, low/low average annual temperature and salinity values. Amplitudes were 4 ^°^C and 2 ppt for all simulations. All simulations began on the coldest, least saline day of the year. Note that increasing temperature increases parasite numbers and decreases generation time. Note also that generation time decreases as the simulation progresses, due to warming temperatures. The sea lice population dies off under the low/low temperature/salinity condition.

**Figure 3:**
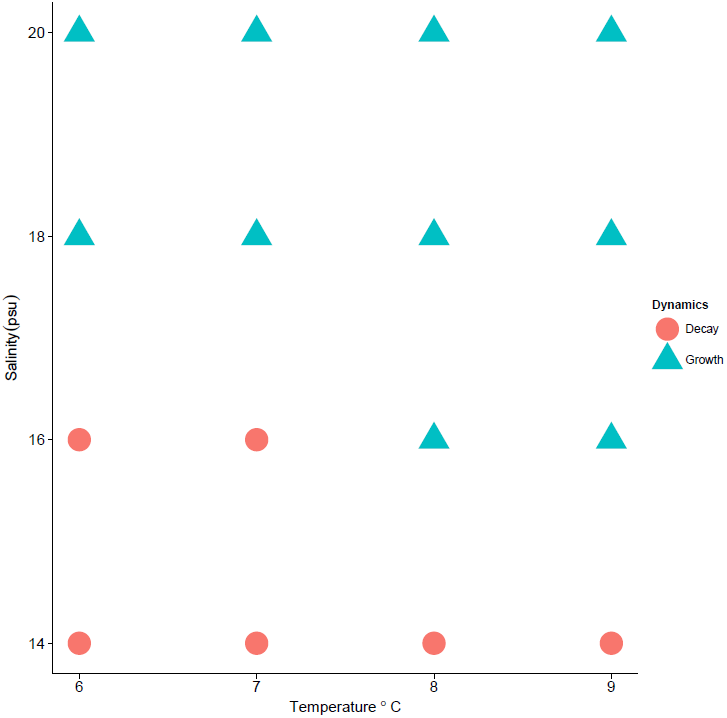
Long term persistence/extinction dynamics at sixteen temperature and salinity combinations. Amplitudes were 4 ^°^C and 2 psu for all simulations. Environmental conditions that result in > 1 adult female sea louse per fish after four years are shown as triangles. Environmental conditions that result in < 1 are shown as circles.

We ran the model with average annual temperatures of 6, 7, 8, or 9^°^C and average annual salinities of 14, 16, 18, or 20 psu. After simulating four years, environmental conditions that resulted in > 1 adult female per fish were considered to be favourable to sea lice, while environmental conditions with < 1 adult female per fish were considered unfavourable. Sea lice in low temperature/low salinity environments will die out. As either temperature or salinity increases, conditions for the sea lice population improve. Sea lice populations are viable at ≥ 18psu at all temperatures investigated and sea lice populations can persist at lower salinities in warmer climates (Fig. 3).

## 5 *R*_0_(*t*)

The basic reproductive ratio, *R*_0_, is commonly used as a measure of reproductive success in populations. The basic reproductive ratio can be defined as the “expected number of secondary individuals produced by an individual in its lifetime” (Heffernen et al. 2005) and acts as a threshold condition that indicates either population persistence or extinction (Caswell 2009). When *R*_0_ ≥ 1 the population will grow with each subsequent generation and persist, whereas when *R*_0_ ≤ 1 the population will shrink with each subsequent generation until extinction. In seasonal systems, *R*_0_ will depend on the time that the infection is introduced to the system. We define *R*_0_(*t*) such that it is the number of second generation adult females produced by a single adult female, who enters the system at time, *t*. As such, *R*_0_(*t*) depends on the probability that the nauplius survives each successive life stage and the timing and duration of egg string hatching during the adult female stage (see Appendix B). We determined *R*_0_(*t*) numerically by augmenting the system of equations (2)–(7) with a delay differential equation describing the number of adult females in the second generation, where this second generation does not reproduce (see Appendix B for details).

In the Broughton Archipelago, temperatures are favourable year round (5th percentile = 6.61, median = 8.90, 95th percentile = 11.75), while salinity levels are very favourable to sea lice survival in the winter and very unfavourable during the summer (5th percentile = 16.11, median = 27.30, 95th percentile = 32.15). The environmental conditions most favourable to sea lice growth are asynchronous because high sea surface temperatures coincide with low salinities and visa versa. We find *R*_0_(*t*) to be highest in December, when salinity is high, but temperatures are low (Fig. 4A). As such, sea lice that enter the farm in December will go on to produce the most offspring despite having longer generation times than sea lice that hatch in the summer months (Fig. 4B). The value of *R*_0_(*t*) is not < 1 at any point during the year, so an adult female that enters at any given time can reasonably be expected to replace itself over the course of its lifetime. *R*_0_(*t*) reaches a peak of 475.13 in December and a low of 23.68 in June and has a mean value of 220.98.

**Figure 4:**
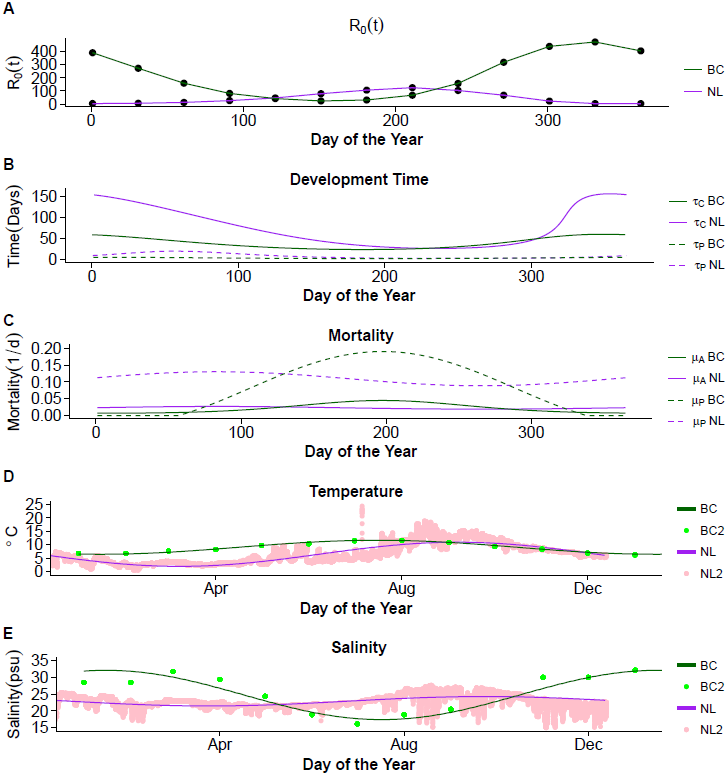
*R*_0_(*t*) is presented for sites in Newfoundland and British Columbia (A). *R*_0_(*t*) is affected by maturation time (B) and the mortality rates of parasitic stages and free-living stages *μ*_*p*_(*t*) (C). Temperature (D) and salinity (E) for the two sites affect development time (B) and mortality (C), respectively.

In southern Newfoundland, temperatures are colder (5th percentile = 2.48, median = 6.00, 95th percentile = 13.20) than the Broughton Archipelago site and sea lice maturation can take a long time during the winter months. Salinity is mostly constant year round (5th percentile = 19.92, median = 22.77, 95th percentile = 26.06), with a low in the spring months. In southern Newfoundland, the environmental conditions that are most favourable to sea lice growth are synchronous: both high temperatures and high salinities occur towards the end of the summer. We find that *R*_0_(*t*) is highest in August (Fig. 4A), after maturation times plummet during the summer months (Fig. 4B) and when time to maturity is shortest (Fig. 4B). The value of *R*_0_(*t*) is not < 1 at any point during the year, so an adult female that enters at any given time can reasonably be expected to replace itself over the course of its lifetime. *R*_0_(*t*) reaches a peak of 125.41 in August, a low of 3.27 in December and has a mean of 46.72.

## 6 Sensitivity Analysis

We conducted a sensitivity analysis on seven model parameters to analyze the effects of their variation on the abundance of female adult sea lice (A). Parameter distributions were estimated from the literature (Table 3). A Latin Hypercube Sampling (LHS) scheme was used to sample the parameter space and partial rank correlation coefficients (PRCC) were used as a test statistic for the sensitivity analysis (Blower & Dowlatabadi 1994).

**Table 3:**
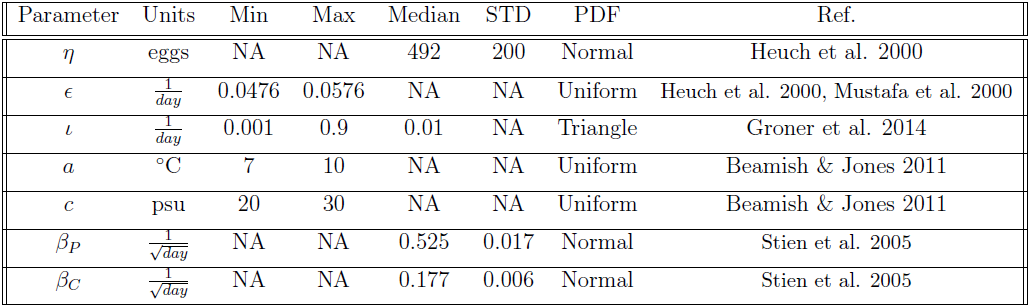
Parameter values for sensitivity analysis

| Parameter | Units | Min | Max | Median | STD | PDF | Ref. |
| --- | --- | --- | --- | --- | --- | --- | --- |
| $\eta$ | eggs | NA | NA | 492 | 200 | Normal | Heuch et al. 2000 |
| $\epsilon$ | $\frac{1}{day}$ | 0.0476 | 0.0576 | NA | NA | Uniform | Heuch et al. 2000, Mustafa et al. 2000 |
| $\iota$ | $\frac{1}{day}$ | 0.001 | 0.9 | 0.01 | NA | Triangle | Groner et al. 2014 |
| $a$ | $^{\circ}\text{C}$ | 7 | 10 | NA | NA | Uniform | Beamish & Jones 2011 |
| $c$ | psu | 20 | 30 | NA | NA | Uniform | Beamish & Jones 2011 |
| $\beta_P$ | $\frac{1}{\sqrt{day}}$ | NA | NA | 0.525 | 0.017 | Normal | Stien et al. 2005 |
| $\beta_C$ | $\frac{1}{\sqrt{day}}$ | NA | NA | 0.177 | 0.006 | Normal | Stien et al. 2005 |

The simulation begins with only adult females and the only parameter that affects adult mortality is mean salinity *(c,* Fig. 5A). After the cohorts start maturing the size of the adult female population is also affected by parameters relating to maturation, infection, and reproduction (Fig. 5A and B). The three most sensitive parameters at 180 days were mean temperature (*a*), mean salinity (*c*), and the number of eggs per egg clutch (*η*; Fig. 5A and B). Female sea lice abundance is more sensitive to the temperature-development relationship of the chalimus and pre-adult stages (*β*_*C*_) than it is to the temperature-development relationship of the nauplius stage (*β*_*P*_; Fig. 5A). Despite a large level of uncertainty about the value of the infection rate, the model is least sensitive to infection rate, out of the seven parameters examined (Fig. 5B).

**Figure 5:**
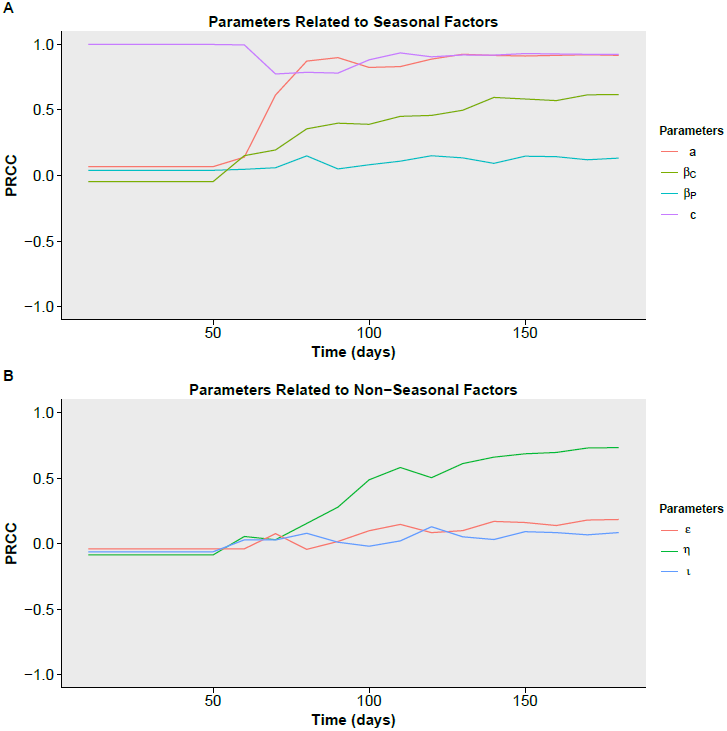
Sensitivity to 7 parameters. The PRCC values at 180 days (from highest PRCC to lowest) are *c* = 0.92, *a* = 0.92, *η* = 0.73, *β*_*α*_ = 0.62, *ε* = 0.18, *β*_*P*_ = 0.13, and *ι* = 0.08.

## 7 Discussion

Sea lice control is one of the top priorities in aquaculture research. Temperature and salinity affect maturation rates, mortality, and egg viability; so control of sea lice relies on understanding their population dynamics in relation to their environment. We derived a deterministic model of the sea louse lifecycle, with temperature-dependent maturation and salinity-dependent mortality. We conducted numerical analyses to: characterize sea lice population dynamics for different environmental conditions, determine the time-dependent basic reproductive number, *R*_*0*_(*t*), for BC and southern Newfoundland, and perform a sensitivity analysis.

There is a substantial difference between the timing of the peak in *R*_0_(*t*) for British Columbia and Newfoundland. We found that the peak value of *R*_0_(*t*) for both the British Columbia and the Newfoundland sites occurred during peak salinity levels, although in Newfoundland the salinity levels were fairly constant and the highest salinity levels also coincided with the highest sea surface temperatures. As such, optimal treatment schemes will also differ between these two sites. In both the Broughton Archipelago in British Columbia and southern Newfoundland, fecund sea lice are able to replace themselves at all times of the year, however, well-timed treatments may result in slower population growth, ultimately leading to fewer required treatments over a production cycle.

We also found that the mean *R*_0_(*t*) was much higher at the British Columbia site than at the Newfoundland site. In considering just the environmental data, it is not clear that this would necessarily be the case. On the one hand, British Columbia has higher mean temperatures, higher mean salinity, and higher maximum salinity: all conditions that are conducive to sea lice growth, while on the other hand, British Columbia also has lower minimum salinity and favourable environmental conditions for sea lice growth are not synchronous as they are in Newfoundland. Contrary to our results, it is generally acknowledged that control of sea lice is typically more straightforward in British Columbia than in Newfoundland. Our model suggests that something other than temperature and salinity may be responsible for the more successful management of sea lice in British Columbia. This may be the result of farm-level and regional-level management decisions, population size of wild hosts, potential genotype differences in sea lice populations (Yazawa et al. 2008; Skern-Mauritzen et al. 2014), or hydrodynamical differences.

The comparison of the British Columbia and the Newfoundland sites suggest no general patterns. To understand the dynamics of sea louse fecundity at other sites, environmental data would need to be provided for analysis using our model. This is especially pertinent as salinity patterns may vary substantially over small spatial scales due to their proximity to rivers, and even two sites within the same broad geographic region potentially could have very different salinity patterns.

It is important to note that the *R*_0_(*t*) we calculate is not a threshold condition for sea lice epidemics, since subsequent generations will hatch throughout the year and experience their own *R*_0_(*t*) values. *R*_0_(*t*) also does not show the size of, or even the instantaneous growth rate of, the population. Rather, our *R*_0_(*t*) provides a means of comparing reproductive output at different times of the year. The methods outlined in Zhao (2015) provide a framework to analytically determine how environmental conditions affect the threshold for sea lice outbreaks. One of the advantages to using a deterministic delay differential equation approach is that the theoretical approaches outlined in Zhao (2015) can be utilized.

Our sensitivity analysis found that adult female sea lice abundance is most sensitive to average annual temperature and salinity. This is likely because a large number of parameters depend on temperature (*τ*_*P*_(*t*) and *τ*_*C*_ (*t*)), salinity (*μ*_*P*_ (*t*), *μ*_*I*_ (*t*), *μ*_*C*_ (*t*), and *μ*_*A*_ (*t*)) or both *(ø_P_* (*t*), *φ*_*C*_ (*t*)). Our findings that lice abundance is more sensitive to the development rate of the combined chalimus/pre-adult stage than to the development rate of the nauplius stage is in line with sensitivity analyses conducted by Revie et al. (2005) and Groner et al. (2014), who found that sea lice numbers were most sensitive to the survival through the pre-adult stage, of which development time plays a major role.

In addition to the need for more empirical studies into the temperature-maturation relationship of chalimus/pre-adult stages that has already been highlighted, a number of avenues exist to improve model accuracy and usability. Because eggs per clutch (*η*) and egg clutch production rate (*ε*) are two parts of the same product, the difference in PRCC values between them is solely due to the distribution we sampled from for each parameter. Egg clutch size is highly variable and the number of eggs per egg string has been suggested to be dependent on the temperature history of the female louse during development (Ritchie et al. 1993; Heuch et al. 2000). Sea lice that develop in colder temperatures are suggested to produce more eggs per egg string as a compensatory strategy for slower development, although the exact mechanism linking temperature history and egg string length is unknown (Ritchie et al. 1993). As such, a better understanding of the relationship between temperature and egg production is needed before it can be incorporated into mechanistic models of sea lice development. Difficulties in mate finding at low densities has also been indicated as a major facet of reproductive success (Stormoen et al. 2013; Groner et al. 2014). Future models may wish to explore the impacts of these biological complexities.

Salmon farms regularly treat for sea lice, which impacts population numbers at a level greater than environmental factors. The timing of treatment, in regards to the typical salmon farming cycle has been shown to have a large impact on sea lice numbers (Revie et al. 2005). Because salmon can be introduced to saltwater pens at most times of the year, our model is well suited to examining the effects of treatment timing on sea lice numbers. We conclude by recommending that future modelling studies incorporate detailed seasonal characterists of their chosen study site into models of sea lice population dynamics.

